# Analyzing Gaze and Hand Movement Patterns in Leader-Follower Interactions During a Time-Continuous Cooperative Manipulation Task

**DOI:** 10.1101/2025.09.29.679248

**Authors:** Minghao Cheng, Anoushiravan Zahedi, Ricarda I. Schubotz, Florentin Wörgötter, Minija Tamosiunaite

**Affiliations:** Inst. Physics 3, Computational Neuroscience, Georg-August University Göttingen, Germany; Faculty of Informatics, Vytautas Magnus University, Kaunas, Lithuania; Department of Psychology, University of Münster, Münster, Germany; Otto-Creutzfeldt-Center of Behavioral and Cognitive Neuroscience, University of Münster, Münster, Germany

**Keywords:** Eye tracking, eye and hand coordination, joint action, predictive coding, Bayesian hierarchical modeling

## Abstract

In daily life, people often interact by taking on leader and follower roles. Unlike laboratory experiments, these interactions unfold naturally and continuously. Although it is well established that gaze typically precedes object manipulation, much less is known about how gaze–hand patterns evolve in interactive settings where one person must take the other’s actions into account. Here we examine predictive, planning-related behavior in a two-player tabletop game called “do-undo.” Participants alternated as Leader and Follower. The Leader performed simple pick-and-place actions to alter the arrangement of objects, while the Follower used other objects to restore the previous configuration. We recorded eye and hand movements, along with object trajectories, using a system that combined eye tracking with multi-camera motion capture. Touch sensors on the players’ hands provided precise timing of contacts, allowing us to segment cooperative action into well-defined temporal intervals. As expected, eye fixations consistently preceded manipulation, but clear role differences emerged. Leaders looked more often and earlier at target objects. In many trials, their gaze anticipated not only their own actions but also those required of the Follower. Leaders also more frequently checked the outcome of the do-undo sequence. Both roles showed gaze patterns consistent with memorization, but alternating gazes between objects and destinations were much more common in Leaders. Some patterns suggested longer-term planning beyond the immediate action. These findings reveal distinct decision-making and planning strategies in Leaders and Followers. Leaders consider not only their own next moves but also the potential actions of their partners, shedding light on the complex cognitive processes that underly everyday human interaction.

## Introduction

Collaborative human-human interaction happens frequently in our daily lives, for example, when we assemble furniture together, fix a bike, or cook a meal together. In these everyday moments of shared action, one person often takes the lead – at least for some duration – and the other follows suit, performing complementary actions to achieve a common goal. Such interactions are often interspersed with intervals where both perform tasks in parallel.

Here, we focus on the leader-follower phases of interaction, as they are cognitively demanding: They involve both agents’ continuous acquisition of information, typically through vision to monitor the other’s actions. This is reflected in gaze behavior, as both closely observe the other’s manipulations. However, eye movements are expected not only to follow an action, but also often to precede it, for example, when looking at a target destination before the object in hand is moved there. A recent transdisciplinary review highlights that eye gaze and visual attention provide reliable yet underused behavioral markers of leadership, followership, and the hierarchical functioning of teams (Cheng et al., 2023). Building on this perspective, recent advances in interpersonal eye-tracking demonstrate that gaze provides a timely method for capturing the real-time dynamics of interacting minds, illuminating how people coordinate attention and build shared understanding during joint action (Wohltjen & Wheatley, 2024).

While we know quite a lot about how individuals coordinate during joint tasks, less is known about the distinct ways leaders and followers use their gaze to predict each other’s movements, especially in object manipulation. How do these anticipatory gazes differ between roles, and how can we measure these differences with enough precision to better understand the underlying cognitive processes?

Research on non-cooperative or individual tasks has consistently shown that gaze precedes manual action. Fixations are often directed to task-relevant targets shortly before the corresponding movement — a “just-in-time” mode of anticipatory gaze (Keshava et al., 2024). At the same time, gaze can also anticipate over longer horizons: for example, it shifts toward the object to be manipulated before the actual action (Land et al., 1999). Furthermore, patterns of repeated fixations on the same object - when the eyes return to it multiple times - have been closely associated with the cognitive processes underlying action planning (Sullivan et al., 2021; Mennie et al., 2007, Pelz and Canosa, 2001).

Clearly, overall gaze patterns are complex, as gaze can also sometimes fall on action-irrelevant objects, or objects may be grasped without a preceding fixation (Hayhoe et al., 2003). The latter might be associated with different visual modalities; for instance, grasps can occur using peripheral vision alone (Brown et al. 2005). It has also been shown that fixations may occur on empty space where an object once was, reflecting memory processes (O’Regan, 1992; Foerster, 2019).

Not only have actors’ fixations been studied, but also those of observers. Similar to actors, observers make predictive looks toward the actor’s manipulated objects (Flanagan and Johansson, 2003), and such predictive fixations are more common than sustained tracking of the actor’s hand (Flanagan et al., 2013). Furthermore, when an observer has previously performed the action themselves, they tend to predict earlier (Möller et al., 2015).

Despite existing knowledge of actors’ and observers’ gaze behavior, much less is known about gaze behavior during joint manipulation actions. For example, several studies examined gaze in cooperative settings involving reference acts (e.g., saying “look at the cup”, or pointing; Gergle and Clark, 2011, Andrist et al., 2015). As the current study does not use verbal communication, gestures, or other referencing cues, we do not discuss this literature further.

Here, we investigate joint tabletop manipulation actions, more specifically hand-object interactions without verbal communication. These interactions remain gaze-intensive because one has to visually plan and attend to one’s own action, while also observing what the other is doing. Existing studies of a similar kind often rely on artificial settings, including sparsely distributed objects of exaggerated size (Huang et al., 2015), virtual reality with large screens (Andrist et al., 2017; Fuchs and Belardinelli, 2021), or specially designed robotic setups for slowing down human motion (Stolzenwald and Mayol-Cuevas, 2018). Although such settings allow improved resolution of eye fixations, they render the environment less ecologically valid. Often, such studies focus on predicting the other’s intentions in a black-box manner, aimed at applications rather than theoretical explanation.

From eye fixation studies of joint actions with more complex tabletop tasks performed in realistic settings, studies show that mutual gaze coordination (i.e., looking at the same location) is much reduced when participants do not communicate verbally (Hessels et al., 2023). In such demanding tasks, participants rarely looked at each other’s faces (less than 0.5% of the time). A follow-up study (Hessels et al., 2024) further investigated the coupling of gaze with one’s own versus the other’s actions. The main finding was that gaze is more strongly coupled to one’s own action. In the same study, speech episodes and gestures were also investigated, yielding similar results. However, looking at individual objects was not differentiated in these studies, which only distinguished larger table regions.

To address these limitations, our study investigates predictive, planning-related gaze behavior in a two-player tabletop game, focusing on the leader-follower dynamic, with the aim of enhancing both the ecological validity of the task and the precision of gaze measurement.

We track eye movements, hand movements, and individual objects on the table and determine how eye and hand movements relate across players. Data were recorded using a setup that combined gaze tracking with multicamera motion tracking of the participants and table configuration, along with touch sensors on the hands. This setup allowed accurate phasing of cooperative manipulation based on the moments of hand contact (*touching*) and release (*untouching*) during object manipulation.

The study investigated whether, in an ecologically valid game where joint action between two players is required, planning and resource allocation would differ when participants assume the role of a leader versus a follower. Further, since leaders must plan actions ahead whereas followers adjust to the leader’s actions, we were interested in the interactions between role (i.e., leader vs. follower), activity (actor vs. observer), and event (planning to grasp an object vs. placing it at a destination). To achieve this, we analyzed the number of fixations, total fixation duration, and fixation latencies using multilevel Bayesian generalized linear modeling.

## Methods

### Subjects

We conducted experiments with a total of 60 adult right-handed participants (39 male, 21 female; age range, 19-35) in pairs. All participants had normal or corrected-to-normal vision. All participants were informed about the purpose of the experiment and gave written informed consent. The experiment was performed in accordance with the ethical standards laid down by the 1964 Declaration of Helsinki and relevant guidelines of the DPG. Furthermore, the experimental procedures were approved by the Ethics Committee of the University of Göttingen, Psychology (Registration Nr: 294).

### Experimental Paradigm

For this study, we designed a paradigm consisting of an Action-Counteraction (ACA) game. The two participants assumed different roles during the ACA game: Leader and Follower. The Leader was free to execute actions, while the Follower responded to the Leader’s preceding action.

At the start, a set of glasses and cubes was placed on the table in two rows with nine locations each. The different allowed object combinations are shown in Figure 1A, and Figure 3A shows the setup on the table. When the game starts, the Leader performs an action, freely chosen from the allowed configurations, including singletons. Upon completion of the Leader’s action, the Follower executed a counteraction. The counteraction had to satisfy two rules: (1) after the pair of action and counteraction, the configurations of the objects on the table must be the same as before. (2) The Follower could not manipulate the same object as the Leader. For example, if the Leader chooses to invert a glass as their action, the Follower must invert another glass back, as illustrated in Figure 1B. Or – more complex – if the Leader un-stacks a glass that is placed on top of a cube and stacks it onto another glass, the Follower must un-stack another glass placed on top of another glass and then place the glass onto another cube in the same orientation (Figure 1C). The setup was designed so that a counteraction is always possible for any of the allowed actions that the Leader can take. To this end, we placed two single cubes and two glasses at 4 locations and filled the remaining ten locations with the allowed object combinations. As a consequence, the four remaining locations remained free. After each action pair, the Leader and the Follower perform the next pair, repeating this for 10 rounds. Then the two players swapped roles and performed 10 more rounds. After 20 rounds, the experimental session ends.

**Figure 1.**
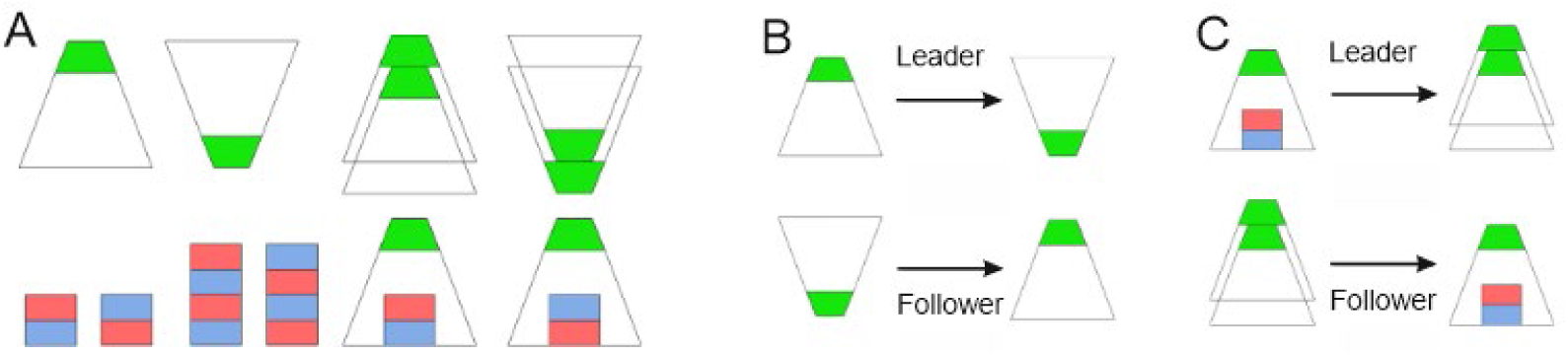
A) Objects used in the ACA game and their allowed configurations. B) Simple example of an action-counteraction pair (glass inverting). C) More complex compound action example.

### Experimental Procedure

Participants sat opposite each other at a round table (see Figure 3A). Each session began with the experimenter explaining the game rules and allowing participants to practice until familiar. Once both participants were ready, they put on white gloves to improve the computer vision system’s recognition of their hands. In addition, the gloves contained a microswitch under the index finger to accurately record touching events. Eye-trackers (Pupil Core eye trackers from Pupil Labs) were attached to the participants, similar to wearing eyeglasses. This was followed by a calibration procedure in which participants look at four different locations on the table. Afterward, the actual recording of the session began. We recorded touching and untouching events using the touch sensors (switches in the gloves) and hand movement patterns using a multi-camera system (see below), as well as the eye movements of both participants simultaneously. During each session, participants engaged in playing the ACA game. Each session consisted of four blocks, each with 20 action–counteraction pairs. In every block, both players assumed both roles (Leader for 10 pairs, Follower for 10 pairs), with order counterbalanced across participants. Once all four game blocks were finished, the session ended. Each block lasted approximately 3 to 5 minutes, resulting in a total session time of less than 20 minutes. In total, 120 data sets were recorded (30 pairs × 4 blocks). Of these, 110 were valid; 10 were excluded due to recording failures.

### Recording Setup

#### Overview

As depicted in Figure 2, the system consists of three input data sources, namely the five-view camera setup from FLIR (Teledyne FLIR LLC), the two Pupil Core eye trackers from Pupil Labs (Kassner et al., 2014), and the touch sensors. They were synchronized using a universal clock, and jointly calibrated using Aruco markers (Apritag 36h11 family, Garrido-Jurado et al., 2014). The touch sensor was designed to capture the touching/untouching events between hands and objects (T/U events).

**Figure 2.**
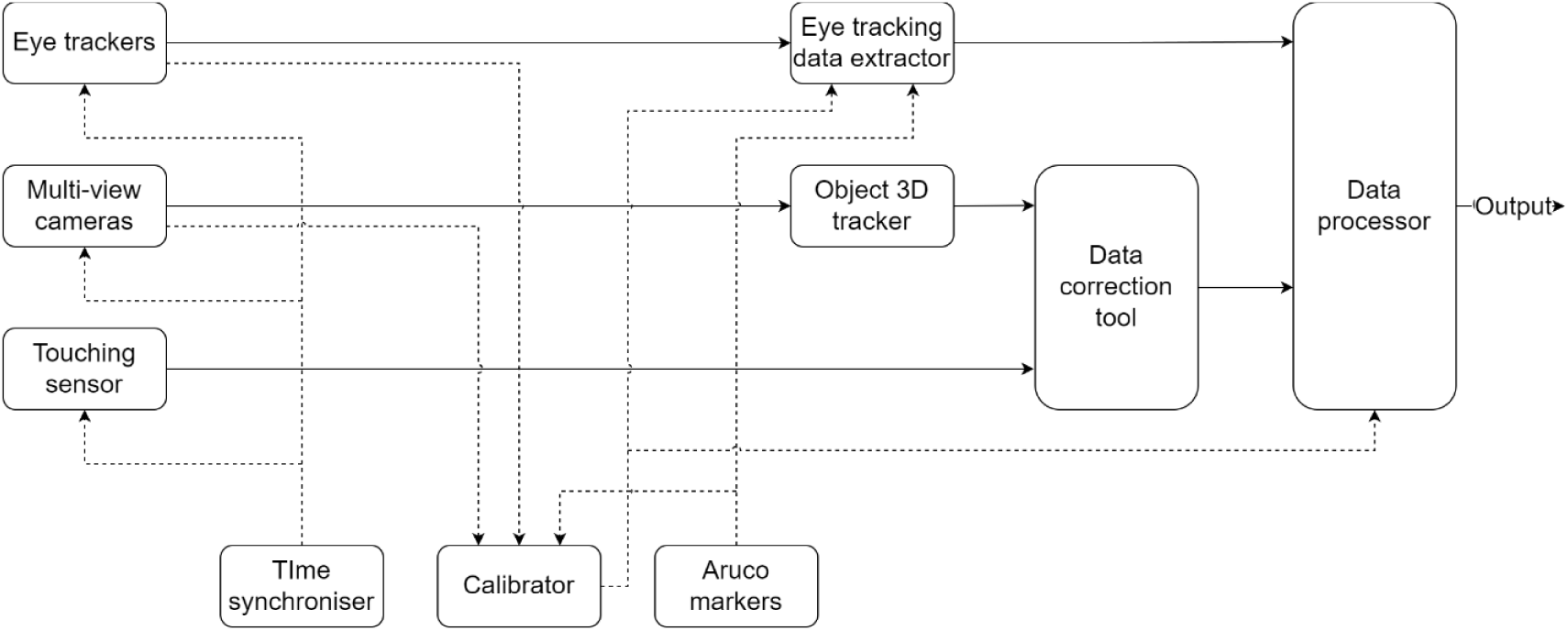
Block diagram of the experiment’s technical setup.

The system’s timing and data flow were as follows. Two calibration procedures were performed. First, a general calibration procedure was conducted. During this process, the experimenter generated a 3D representation of the tablecloth filled with Aruco markers using the Pupil Labs eye tracker and associated recording software. Subsequently, a series of images of the same tablecloth was captured by the five-view camera setup, allowing for the generation of a corresponding 3D model. The calibration software was then used to compute the transformation matrix between the five-view camera configuration and the eye trackers. This calibration procedure only had to be done once unless there are alterations in the physical positioning or orientation of the table or cameras. Second, for each experiment, the experimenter calibrated the eye trackers for both participants as described above, and then triggered the start of the recording. The incoming video stream from the five-view camera setup was encoded and written to the hard drive in real-time, alongside the data acquired from the eye trackers and the touch sensors. The recorded data were analyzed offline. During this process, an object tracker identified and tracked objects and hands, and also extracted the head direction present in each video frame. Finally, the data of interest were extracted and analyzed.

### Hand tracking

The hands of the participants were tracked using an Axis-Aligned Bounding Box (AABB) pipeline. Using the recorded camera images from all five cameras, a custom-trained Deep Neural Network (DNN) -YOLOV5 model (Jocher et al., 2020) detected hands in the images and outputs 2D bounding boxes. Subsequently, a triangulation algorithm transformed these 2D bounding boxes into 3D AABBs. Next, the 3D AABBs were tracked using a modified Unscented Kalman Filter (UKF) and finally smoothed with a low-pass Finite Impulse Response (FIR) filter. See Figure 3A for a view of the scene that includes a 3D AABB on one of the objects.

**Figure 3.**
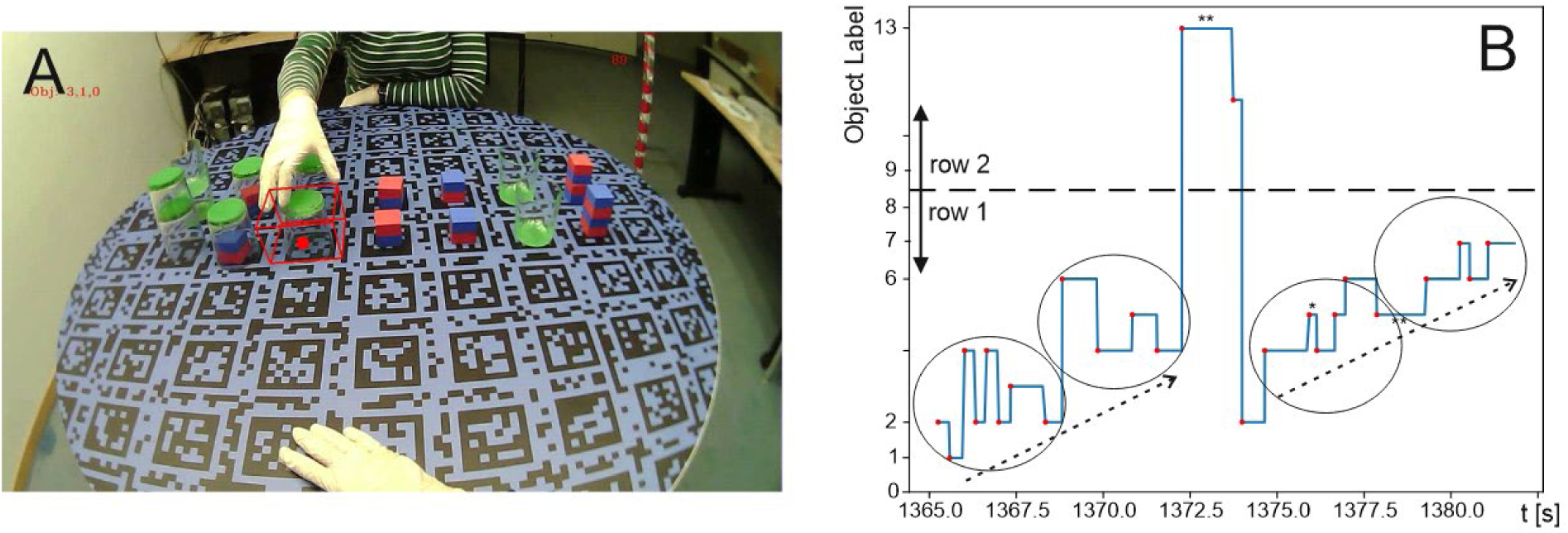
A) View as seen by the camera of the eye-tracker with object bounding box and gaze-indicator (red dot) of the person in front included. This view is essentially identical to the way participants see the scenery. B) Eye track over time. The start of fixations is marked by red dots.

### Object detection

Object detection and label assignment used the same Yolov5 model. The labels of the different objects were assigned to the various possible object positions (see below) after the completion of a manipulation action.

### Eye-tracking data extraction

In this study, the eye tracking data of each player were defined as an array of eye fixations over time. Each array entry contained: the starting time of the fixation; the duration of the fixation; the coordinates of the target position, or the instance ID of the target or hand the participant looked at.

Each eye tracker featured three mounted cameras: one scene camera and two eye cameras. Using the recording software provided by Pupil Labs, the eye tracker output the gaze in the form of 2D coordinates on the scene camera and videos from all the cameras. Two types of data were extracted offline after recordings using the software provided by Pupil Labs. One type was the fixation data, which is defined as a set of consecutive frames where the gaze remains at the same point for a sufficiently long period. To determine a fixation, a threshold of +/-2.45 degrees of visual angle was set within which gazes were attributed to the same object. Furthermore, a resolution of 20ms for fixation shifts was chosen such that the initial sampling rate of the eye-movement data was 50Hz. A fixation was defined when at least 10 such samples hit the same object; hence, we assumed a minimal fixation duration of 200ms (Johansson et al., 2001, Pannasch et al., 2008). Sometimes, glitches occurred where samples from the eye-tracker had jumped away from the actual target object, and these were filtered out.

The second type of data extracted by the Pupil Player software was the head pose data. By detecting the Aruco markers on the table and employing the Perspective-n-Point (PnP) algorithm, we determined the extrinsic parameters of the eye-tracker’s scene camera. With the head pose data, the 2D gaze fixation was transformed into 3D, which was essential for integrating eye tracking data with the location of the objects.

### Determining hand and object location

To calculate which hand or object location the eye fixations struck, a ray-tracing algorithm based on the principles outlined in Williams et al. (2005) was implemented in this study (see Appendix). Ray tracing was essentially a collision detection algorithm between a virtual ray originating from the eye and extending towards the object. This determined which object was being fixated.

To conveniently describe the positions of the objects as well as empty target positions, a discretized coordinate system of the table and a set of 18 corresponding virtual 3D AABBs was defined. This was possible because no other object locations were permitted in this game. In Figure 3A, the defined positions were based on the two rows of Aruco markers on the table. The top left corner of the marker corresponds to the defined location with coordinates (0, 0), and the bottom right corner corresponds to (8, 1). Notably, z-coordinates were not required for the present analyses. There were two reasons that virtual 3D AABBs were used in this study. First, they were used to detect whether the participants looked at empty positions on the table. Second, since the objects were relatively small in size, using the virtual 3D AABBs allowed more precise determination of which object the participants looked at, as the objects could only be placed on these defined, discretized positions. Note that object labels could be assigned to the different locations following object detection as described above.

Figure 3B illustrates an exemplary eye-movement track from one of the sessions, demonstrating the basic data structure used for all statistical analyses. As mentioned, objects are arranged in rows 1 and 2 and numbered from 0 to 8 (left to right) in row 1 and from 9 to 17 in row 2, with the diagram truncated above 13 because no saccades occurred to objects 14–17. For this participant, row 1 lay directly in front, and row 2 was further back. The track shown here contains a total of 25 saccades to different objects over a period of 17.5 seconds. Minimal fixation duration was 220ms (marked by * near 1375.0 s in the figure) and maximal duration 1500ms (** at and after 1372.5 s). Clearly visible are two progressive sequences of saccades along row 1 (dashed arrows), interrupted by looking at row 2 for a short time. During these progressions, alternating saccades to neighboring (or close-by) objects, highlighted by the ellipses, are found.

### Data Analysis

Data were analyzed in two ways: 1) with a time-resolved approach and 2) based on statistical modeling.

### Methods for time-resolved analysis

The raw eye- and hand-tracking data were analyzed with several methods. For hands we determined two data points: 1) the start of the hand movement, determined from the camera images as the moment when the hand leaves the “home” position for more than 5cm and 2) the moments when the hand touches (T-event) or releases an object (U-event) as well as the durations in-between, determined from the sensor data.

<u>Chunking:</u> This resulted in a natural chunking of the data stream. In Figure 4, a schematic representation of the timed data, these intervals are marked with green and orange bars. Note that, naturally, T-U-intervals alternate between Leader and Follower, and there were intervals in-between, called *object-static intervals*, which represent those periods when no object was moved (although hands could still move). For example, note that the period where the Leader does not move an object stretches from the end of interval 2 to the beginning of 6 (for Follower: end of 4 to beginning of 8), always bridging three intervals. The short green and orange arrows indicate this “bridge”. This repeated for other groups of intervals in the same manner (e.g., 1→3, 7→9, etc.). The longer arrows refer to the bridge that reaches from untouching to the next untouching, and this longer interval was also used (see statistical methods).

**Figure 4.**
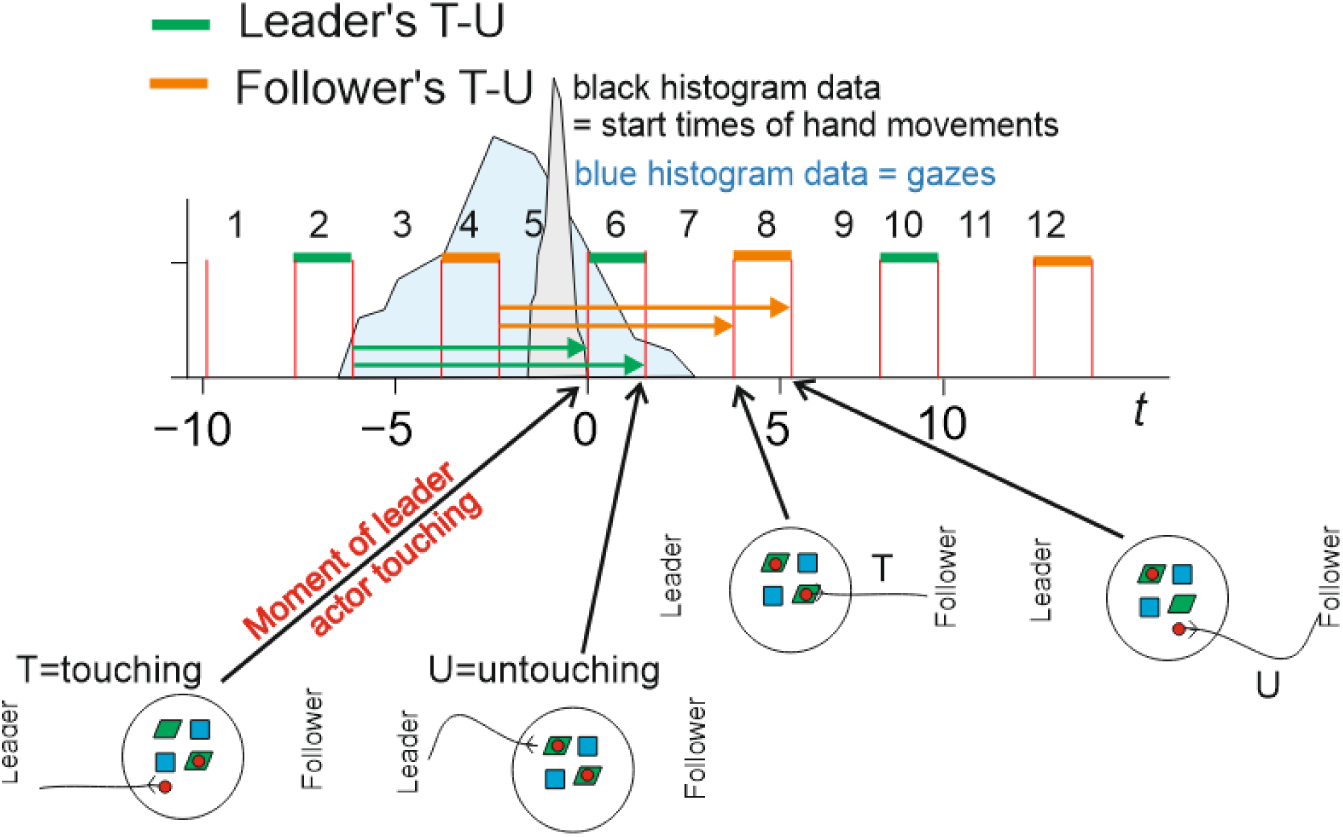
Schematic of the way temporal data will be presented. Arrows refer to the statistical search zones for Leader (green) and Follower (orange) from untouching to next touching (shorter) and from untouching to next untouching (longer).

A total of 12 intervals are plotted in all the Figures that show histograms. This number was chosen to include a total of 3 T-U intervals, each for the Leader as well as the Follower, plus one more object- static interval in the beginning to create a full 3-interval bridge (for the Follower) at the start of each diagram. In the following, we refer to the temporal duration from 1 to 12 as an “episode”.

The pictograms at the bottom show the course of action schematically, where the Leader placed a red disk onto a green parallelogram and the Follower subsequently performed the respective undo operation elsewhere on the table.

We used such a long stretch of an episode lasting from interval 1 to 12 to allow for an extended analysis of viewing behavior, but here care had been taken that during an episode no object was manipulated twice^1^. Episodes where this happened were excluded from the analysis, because, as soon as the same object, let us say O1, was manipulated twice in one episode, it was not clear to which of the two manipulations the fixations on O1 should be attributed. As a consequence, the number of episodes in different histograms is not the same. To account for this, and to make the data comparable, we normalized all histograms to 1000 episodes.

<u>Histogram data representation and its time origin:</u> In addition to the chunking from 1 to 12, Figure 4 also illustrates other aspects. Here, zero of the time-axis refers to the moment of the Leader’s touch of the to-be-manipulated object (start of interval 6). This centering was applied because most gazes were expected – and indeed observed (see below) - before this moment in time. This is shown in the histograms below, where the blue schematic represents this alignment. The numerical values on the abscissa represent, accordingly, the average times before or after in seconds. The same zero-centering was also used for the Follower and, their time zero referred to the start of interval 8 (see Figure 5 and Figure 6). In black, we encode histograms for the starting moments of the hand movements (see panels A and C in Figure 5 and panels A and E in Figure 6).

**Figure 5.**
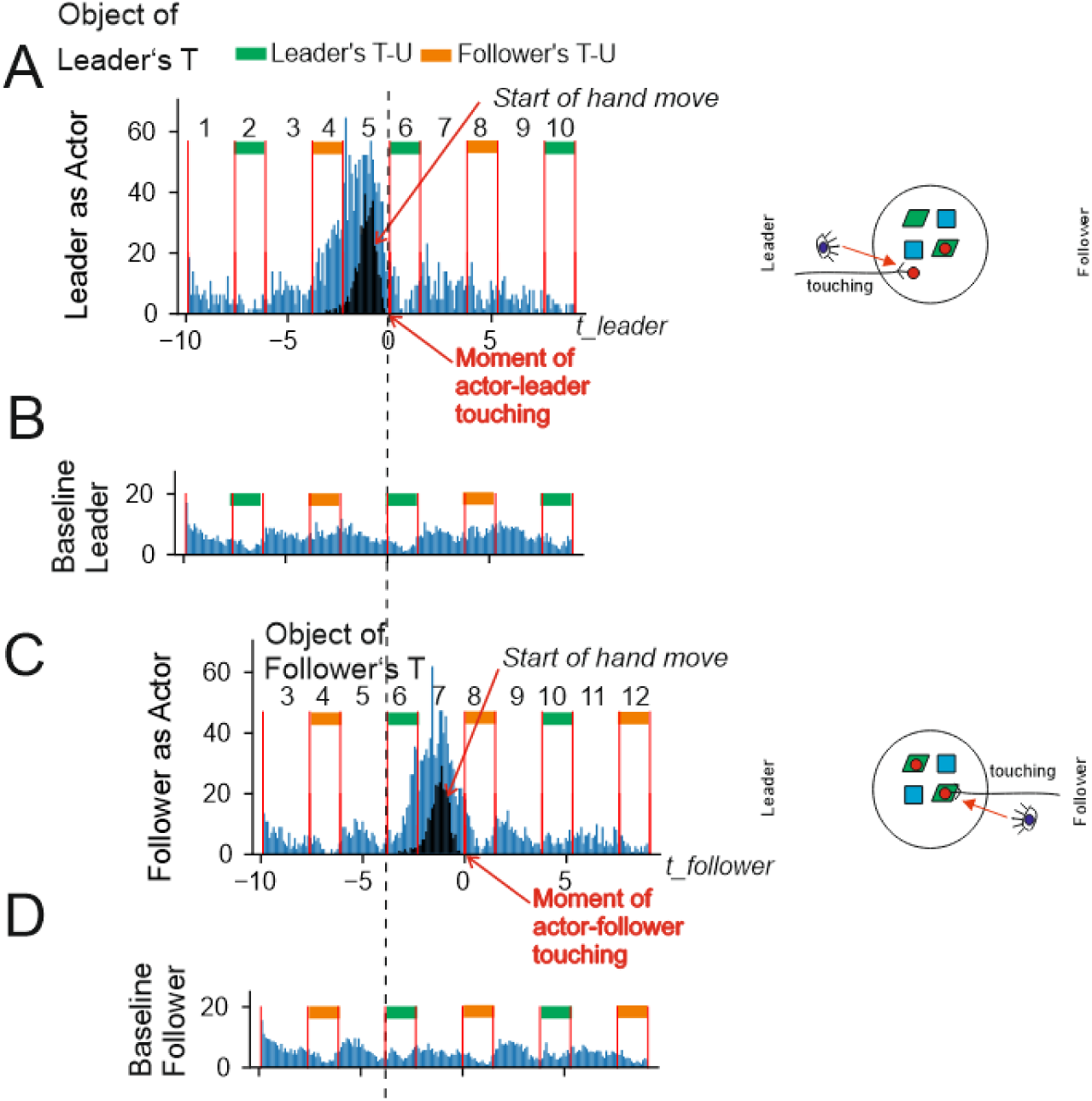
Histograms for two situations and corresponding baselines. Here and in the following: when a diagram is labeled "Object" (panels A,C) it represents the gazes at that particular object that is being manipulated during this episode and similarly for “Destination” (see next Figure) of where that object was put down. Histograms are calculated to represent the number of fixations onto object or destination over all participants and all objects/destinations, where the ordinate is normalized by the number of episodes analyzed (see Method section). On the right side, a schematic shows the table configuration looks like at the given moment in time and also what happens for Leader and Follower (touching an object, untouching the object after placing it at a new destination). Hence, panel (A) left side shows the situation where the Leader touches an object (pictogram of the small red disk) and the histogram(s) show the start of the Leader’s hand movement (black) and – as indicated by the eye pictogram – the statistics of the Leader’s gazes at this object (blue). Diagrams are labeled by different action- as well as inactivity-intervals (numbers and orange, green markers).

**Figure 6.**
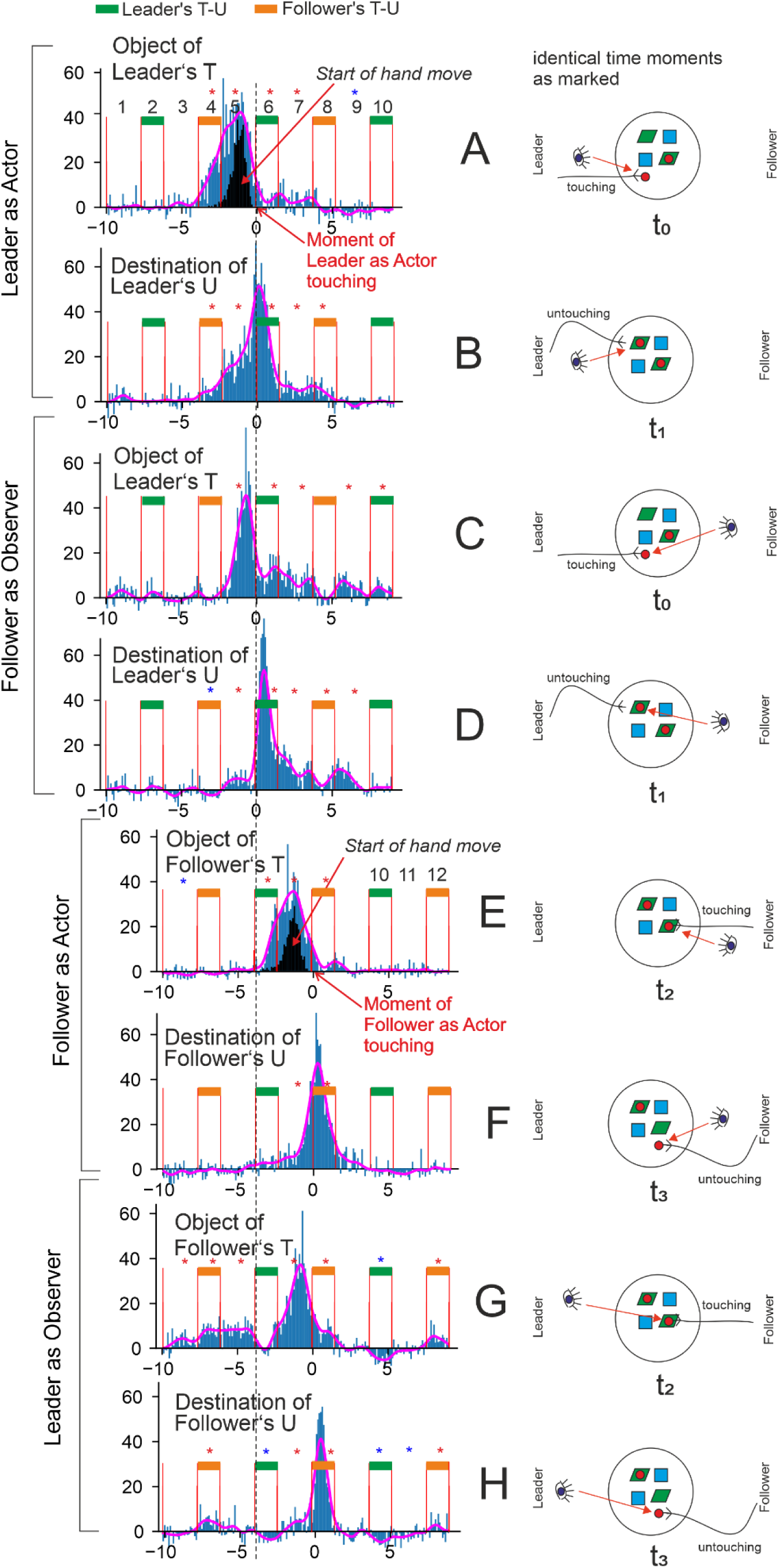
Detailed temporal analysis of hand and gaze patterns. A-H) Actor-Observer, Leader-Follower, and Object-Destination combinations as indicated.

<u>Temporal Normalization:</u> Given that the duration of the intervals was not constant but varied from episode to episode and between different players, we normalized all interval durations in the histograms (Figure 5 and Figure 6) to their respective averages, which were 1.5 s for T–U intervals and 2.3 s for object-static intervals (Leader and Follower alike), and we performed time-warping of all the temporal gaze data within each interval. In a free-running (no triggers, no time limits) experiment, such as the one presented, players were sometimes inattentive or distracted, or for other reasons, overly long or short interval durations would occur. Hence, extreme outliers were removed [*Quartile* 1(*Q*1) − 3 ∗ *interquartile*(*IQR*), *Q*3 + 3 ∗ *IQR*]. We found that about 5% of of intervals fell into this category.

### Statistical Methods

The data were processed and analyzed using the R programming language (http://www.R-project.org/). First, search zones were defined for extracting the relevant gazes for each event (arrows in Figure 4). For Touching and Untouching events, the search zone started from the event and extended until the previous Untouching of the same kind (e.g., the Touching of an observer to the previous Untouching event of the observer). Additionally, filter zones were defined that included the two previous Untouching events of the same kind. To ensure interpretability, all events with two similar target gazes in the filter zone were excluded from the final analyses. This procedure resulted in the elimination of approximately 25% of the events.

The selection of search and filter zones was based on the fact that participants could start planning an action (i.e., grabbing an object and moving it to the chosen destination) only after completion of the previous action, i.e., the previous Untouching. However, as soon as they started planning the action, they could simultaneously plan which object they wanted to move and where they wanted to place it. Therefore, the zones started from the previous Untouching event for both the Touching and Untouching conditions. Furthermore, as planning could only take place before an event, for Touching, the search and filter zones could only extend from the previous Untouching event to the Touching event; however, for Untouching, they would extend from the previous Untouching event to the next Untouching event. To control for the effects of longer search zones for Untouching compared to Touching events, we added the duration of search zones to our models as a nuisance regressor.

For each event, three variables were calculated. (1) The number of correct fixations, defined as the number of matches between the current fixation and the target destination for the event in the search zone. Specifically, the correct fixation for a Touching event was a gaze directed to the location where the object to be moved was located. The correct fixation for an Untouching event was a gaze at the final destination of the object to be moved. (2) The length of correct fixation, defined as the cumulative duration of correct fixations for each event. Finally, (3) the fixation latency, defined as the time interval between a correct fixation and the end of the event. Notably, for each event, several fixation latencies could be extracted, in contrast to the count and length variables, which only had one value per event.

Each event had three categorical properties that were modulated within-subjects: (1) Activity: Actor vs Observer, (2) Role: Leader vs Follower, and (3) Event: Touching vs Untouching. The response variable for each cell of the design was checked for any extreme outliers in the same way as above [i.e., outside *Quartile* 1(*Q*1) − 3 ∗ *interquartile*(*IQR*), 𝑄3 + 3 ∗ *IQR*]. Since the response variables were not modeled under the assumption of a normal distribution, outliers, when present, were not excluded; instead, other techniques, such as using robust statistics and weighted models, were employed to mitigate their effects (Agresti 2015, Huber and Ronchetti, 2011) and are described in the text. Bayesian generalized hierarchical linear regression models (BGLM) were used to investigate each of the variables. For each response variable, the three explanatory variables and their interactions were included in the models. Based on recent developments in statistics (Rouder et al., 2022; van den Bergh et al., 2023), we employed the maximum random-effects model. Thus, we assumed random intercepts and slopes for all included main and interaction effects for each participant (i.e., *sub_id_* in Eq. 1). Finally, to control for differences in search zone length across various events, a search zone length variable was added to the model as a nuisance regressor.

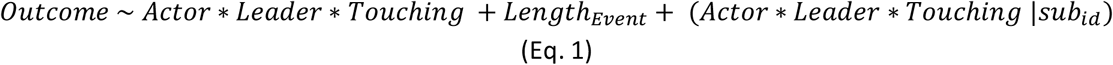

For calculating Bayesian hierarchical generalized linear models, the packages brms (Bürkner, 2017) and RStan (https://mc-stan.org/) were employed. All models were estimated using five chains, each with 4000 iterations and 2000 warm-up iterations. If any variable showed R, the potential scale reduction factor on split chains, above 1.05, the model was recalculated with increased iterations, and the results were reported accordingly. Finally, since all the models were hierarchical, weakly informative priors were preferred (Rouder et al., 2022; van den Bergh et al., 2023). The exact weakly informative priors used for each model are described below.

For modeling the fixation counts (Eq. 2), the response variable represented the number of events occurring within a specific time window. Hence, a negative binomial BGLM was employed. A negative binomial distribution was preferred over a Poisson distribution due to overdispersion. The weakly informative priors for the model were as follows: *N*(0,10) for the intercept, *N*(0,2) for the β coefficients, *student*_*t*(3, 0, 2.5) for the σ and SD hyperparameters, *inv* − *Gamma*(0.01, 0.01) for the shape hyperparameter, and *lkj*(1) for the correlations between random variables.

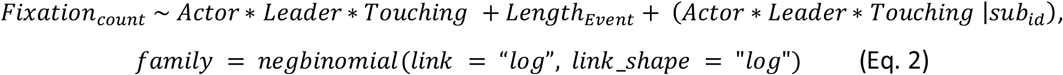

For modeling fixation length (Eq. 3), a hurdle-lognormal model was used (Cragg, 1971; Neelon et al., 2010). The hurdle model allows us to handle zeros (i.e., when no correct gaze was found and thus the gaze length was zero) simultaneously with the rest of the data (lognormal data related to gaze length when there was at least one correct gaze). The weakly informative priors in the models were: (0,10)for the intercept, *N*(0,2) for the β coefficients, *student*_*t*(3, 0, 2.5) for the σ and SD hyperparameters, and *lkj*(1) for the correlations between random variables.

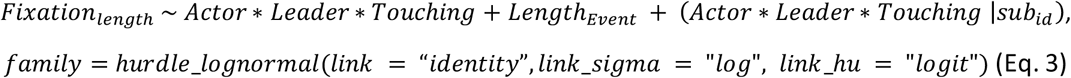

Finally, we used a lognormal model to investigate fixation latency. The weakly informative priors employed for this model were: (0,10) for the intercept, *N*(0,2) for the β coefficients, *student*_*t*(3, 0, 2.5) for the σ and SD hyperparameters, and *lkj*(1) for the correlations between random variables.

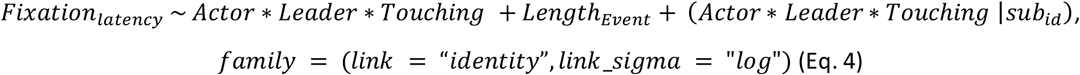

For the model comparison, we used the Pareto smoothed importance sampling (PSIS) estimation of leave-one-out cross-validation (LOO) implemented in the LOO package (Magnusson et al., 2020; Vehtari et al., 2016). LOO assesses pointwise out-of-sample prediction accuracy from a fitted Bayesian model using the log-likelihood evaluated at the posterior simulations of the parameter values; however, as it was difficult to calculate, importance weights ware commonly used instead, resulting in PSIS-LOO. To make sure that PSIS-LOO estimation was accurate, we used *Pareto* 𝑘 < 0.7. However, *Pareto* 𝑘 < 1 was still considered acceptable.

All hypotheses were tested using the function in brms (Bürkner, 2017). Based on the suggestion of van Doorn et al. (2021), *Bayes factors*(*BF*) > 3 were considered as significant evidence for the tested hypothesis. One-sided hypotheses (*BF*_+0_and *BF*_0+_) compared the posterior probability of a hypothesis against its alternative. On the other hand, two-sided tests (*BF*_10_and *BF*_01_) compared hypotheses with their alternatives using the Savage-Dickey density ratio method (Bürkner, 2017).

### Validity

In all cases, the different model variants converged correctly without any divergent transitions (all *Rhat* = 1). Furthermore, the bulk and tail effective sample sizes for main effects and interactions were each above 4000, indicating that model predictions were reliable. The model accurately captured the observed data distribution.

## Results

All analyses used a minimal fixation duration of 200ms, which is realistic threshold in this context (Johansson et al., 2001, Pannasch et al., 2008). Note that we annotate with “object” the grid location of an object about to be manipulated by the actor. We use the same annotation “object” for grid locations from which an object has been removed. With “destination,” we annotate the place where an object was or would be placed. The latter includes all possible places on the game grid (not only empty ones).

First, we provide a time-resolved analysis of general effects, followed by detailed statistical analyses to consolidate the main findings.

### Temporal characteristics of the viewing behavior

Figure 5 shows the viewing behavior as blue histograms alongside hand-movement onsets (panels A, C, black histogram). Histograms were centered (“zero”) at the moment where the Leader (A,B) or the Follower (C,D) touched the object, which is to be manipulated. This was chosen because most gazes were expected - and indeed observed - before this moment as shown by the main blue peaks. Detailed evaluations are provided below (Figure 6). The dashed vertical line visually aligns the top and bottom panels.

When the Leader was the actor (A), they *started* their hand movements during interval 5 (black histogram) and the hand then eventually touches the object. The blue histogram shows the distribution of the Leader gazes at the object being manipulated at this stage. The bulk of this occurred before interval 6. Hence, as expected, eye movements predict hand movements. In addition, there was a tail that extended into intervals 7-9, which will be discussed later. Panel (B) shows a baseline. It contains all gazes of the participants at objects not having been manipulated during the length of the episodes contained in panel (A). Below we used the baseline to calculate statistical significances. Panel (C) shows the corresponding situation when the Follower was the actor, with the baseline given in (D).

Hence, in summary, both histograms A and C represent the gazing behavior at the to-be-manipulated objects, illustrating part of the do–undo sequence: specifically, the object-centered viewing behavior of the Leader who prepares and performs an action and the (also object-centered) viewing behavior of the Follower who then acted to perform the undo action. In Figure 6, we now discuss these and more cases in more detail, showing which intervals display a significant deviation from baseline and which did not. We thus subtracted the baselines from their corresponding original plots, which could lead to negative numbers, too. Red (blue) asterisks represent intervals that were significantly greater (smaller) than baseline level (p<0.01 by a one-sided Wilcoxson signed rank test).

Pink curves represent a low-pass filtered version of the blue histograms, using a Gaussian filter with a standard deviation (STD) of 0.3 s. Note that diagrams come in pairs occurring at the same time (t_0_, t_1_, t_2_,t_3_, indicated under the pictograms). The vertical dashed line marks the start of interval 6, which is the one where the Leader actually starts their action. All four top diagrams (A-D) show a tail behind the main peak. These diagrams are all about the Leader’s object (taking and placing). These tails represent looks “into the past”, either at the place where the object was located before manipulation and/or at the destination previously covered by the object.

We now provide results in more detail for the different figure panels in Figure 6.

Panel (A): This panel illustrates the behavior of the Leader when considering the to-be-manipulated object. It also includes the start of the Leader’s hand movement, which occurred in interval 5 before the touching of the object. The Leader’s gaze shifted to the object just before interval 4, and mainly during intervals 4 and 5. Meanwhile, in interval 4, the Follower continued their task, undoing the previous game event – an action that does not require the Leader’s attention. Intervals 6 and 7 displayed a tail. Intriguingly, at/after their untouch, some Leaders looked again at the location where the object had been taken away from.

Panel (B): This panel illustrates the behavior of the Leader when considering the destination for placing the object. The Leader looked at the destination very often before touching their object (earliest start of looks at end of interval 3). The whole plot was shifted by one interval relative to panel A, which is expected, because targeting a destination must come after targeting an object. Again, a tail was present, lasting until interval 8.

Panel (C): This panel illustrates the behavior of the Follower who observed the object that is later selected by the Leader. Naturally, the Follower looked at the Leader’s object *only* when or after the Leader actually had started to move the hand (compare peak in (C) to the black histogram in (A)). There was only a short delay of approximately 400ms between the peak in panel C and the black peak in panel A. Again, there was a tail of looks at the location where the Leader’s object had been removed from.

Panel (D): This panel illustrates the behavior of the Follower who observed the destination where the Leader placed the object. This observation closely followed interval 6 — the Leader’s action interval — and showed little predictive component. The bulk of the Follower’s gazes occurred just slightly before the Leader’s untouch, and some gazes occurred afterwards. Again, there was a clear longer-lasting after-look tail.

Intriguingly, for diagrams E-H (Follower as actor and Leader as observer) there was no tail after the main peak. Note that at time t3 the initial situation on the table was recovered [compare panels (A,F)].

Panel (E): This panel illustrates the behavior of the Follower when considering the object to be used for the undo action. This diagram resembles a shifted copy of panel A with very much the same characteristics but without the tail. This general shape was expected as here the Follower was preparing to touch their object. Remarkably, when overlaying the smoothed curve of E onto panel A, one finds an almost perfect fit to the main peak in A.

Panel (F): This panel illustrates the behavior of the Follower when considering the destination for placing the object. The same observation as for panel E also held for panel F: It was essentially a shifted copy of panel B without tail. Here, too, the smoothed curve matched the one from above exceedingly well.

Panel (G): This panel illustrates the behavior of the Leader who observed the object that the Follower will take. It shows a somewhat broader peak when looking at the Follower’s object, compared to the Follower’s peak in panel C when looking at the Leader’s object. The main peak occurred before the object is touched by the Follower. By aligning it with the hand movement plot in panel E, it is evident that the Leader uses the hand movement of the Follower to predict which object the Follower will touch. This is again similar to the observation in panel C. However, the main peak in (G) is slightly broader before its maximum compared to the peak in panel C, because the Follower in panel C had no advance information about the Leader’s next action and could only predict based on the Leader’s hand movement. In contrast, the Leader knew that the Follower had only a limited set of possible objects to choose from, which sometimes allowed for an earlier prediction. Furthermore, in intervals 4 and 5 there was a small but significant peak, indicating that – while the Leader was planning their own action (and at the same time observing what the Follower was doing) – they also looked at potential objects the Follower could use in the future to undo their currently planned actions.

Panel (H): This panel illustrates the behavior of the Leader who observed the destination where the Follower placed the object. The histogram was narrow and, quite similar to (D), it began with a slight predictive component during the Follower’s movement (interval 6, orange) and ended with the Follower’s release of the object. A narrow histogram was expected, because, similar to (D), this pattern mainly reflected reactive (non-predictive) observation, occurring sharply in alignment with the Follower’s movement interval (orange).

After exploring the eye and hand movement data, we next applied Bayesian generalized hierarchical modeling to statistically test the differences between different roles, activities, and event types in fixation count, duration, and latencies.

### Fixation count – Number of correct fixations, their distribution, and fixation patterns

#### Fixation count

We used Equation 2 to determine the number of eye fixations on objects and destinations during the task and under different conditions. We studied the main effects and interactions via hypothesis testing (for full results, see Table 1 in the Supplement).

**Table 1.** Hierarchical Bayesian generalized linear modelling results for the fixation count (Eq. 2 in the main text)

| Hypothesis | Estimate (SE) | 95% CI | BF<sub>01</sub> | Post Prob |
| --- | --- | --- | --- | --- |
| Activity = 0 | 0.11 ± 0.04 | [0.02, 0.19] | 1.89 | 0.65 |
| Role = 0 | 0.03 ± 0.04 | [-0.05, 0.11] | 38.25 | 0.97 |
| Event = 0 | -0.05 ± 0.03 | [-0.12, 0.02] | 23.67 | 0.96 |
| **Time Window = 0** | 0.19 ± 0.01 | [0.17, 0.21] | **0.00** | **0.00** |
| **Activity \* Role = 0** | 0.21 ± 0.05 | [0.10, 0.31] | **0.01** | **0.01** |
| Activity * Event = 0 | 0.04 ± 0.05 | [-0.06, 0.13] | **30.58** | **0.97** |
| **Role \* Event = 0** | 0.19 ± 0.05 | [0.10, 0.28] | **0.00** | **0.00** |
| **Activity \* Role \* Event = 0** | -0.23 ± 0.06 | [-0.36, -0.11] | **0.06** | **0.05** |

Note: Significant alternative hypotheses are indicated in bold.
CI: Credential interval.
BF<sub>01</sub>: Bayes Factor comparing the Null hypothesis to the alternative hypothesis. All hypotheses were tested using the brms package (Bürkner 2017). Based on the suggestion of van Doorn et al. (2021), *Bayes factors*(BF) > 3 were considered as significant evidence for the null hypothesis and *BF* < 1/3 for the alternative hypothesis.

The results showed that the interaction between Role and Event was significant (*H_0_: Role*Event* = 0, *Estimate ± SE* = 0.21 ± 0.05, CI = [0.10, 0.31], ***p.p.* = 0.01, *BF_01_* = 0.01**). When being a Leader (*mean*_Untouching−Touching_ = 0.35 ± 0.38) compared to a Follower (*mean*_Untouching−Touching_ = 0.33 ± 0.39), the difference in fixation counts between Touching and Untouching events was bigger.

Additionally, the significant interaction between Role and Activity (*H_0_: Role*Activity* = 0, *Estimate ± SE* = 0.19 ± 0.05, *CI*= [0.10, 0.28], ***p.p.* = 0.00, *BF_01_* =0.00**) revealed that Leaders (*mean*_Actor−Observer_ = 0.34 ± 0.01), compared to Followers (*mean*_Actor−Observer_ = 0.15 ± 0.01), had a higher fixation count when acting compared to observing.

Finally, also the three-way interaction between Role, Activity, and Event was significant (*H_0_: Role*Activity*Event* = 0, *Estimate ± SE* = -0.23 ± 0.06, *CI*= [-0.36, -0.11], ***p.p.* = 0.05, *BF_01_* =0.06**), indicating that although Leaders compared to Followers looked significantly more frequently at objects moved by themselves or their partners, they looked at the destinations significantly more frequently only when acting themselves, but not when their partner was moving an object (Figure 7A).

**Figure 7.**
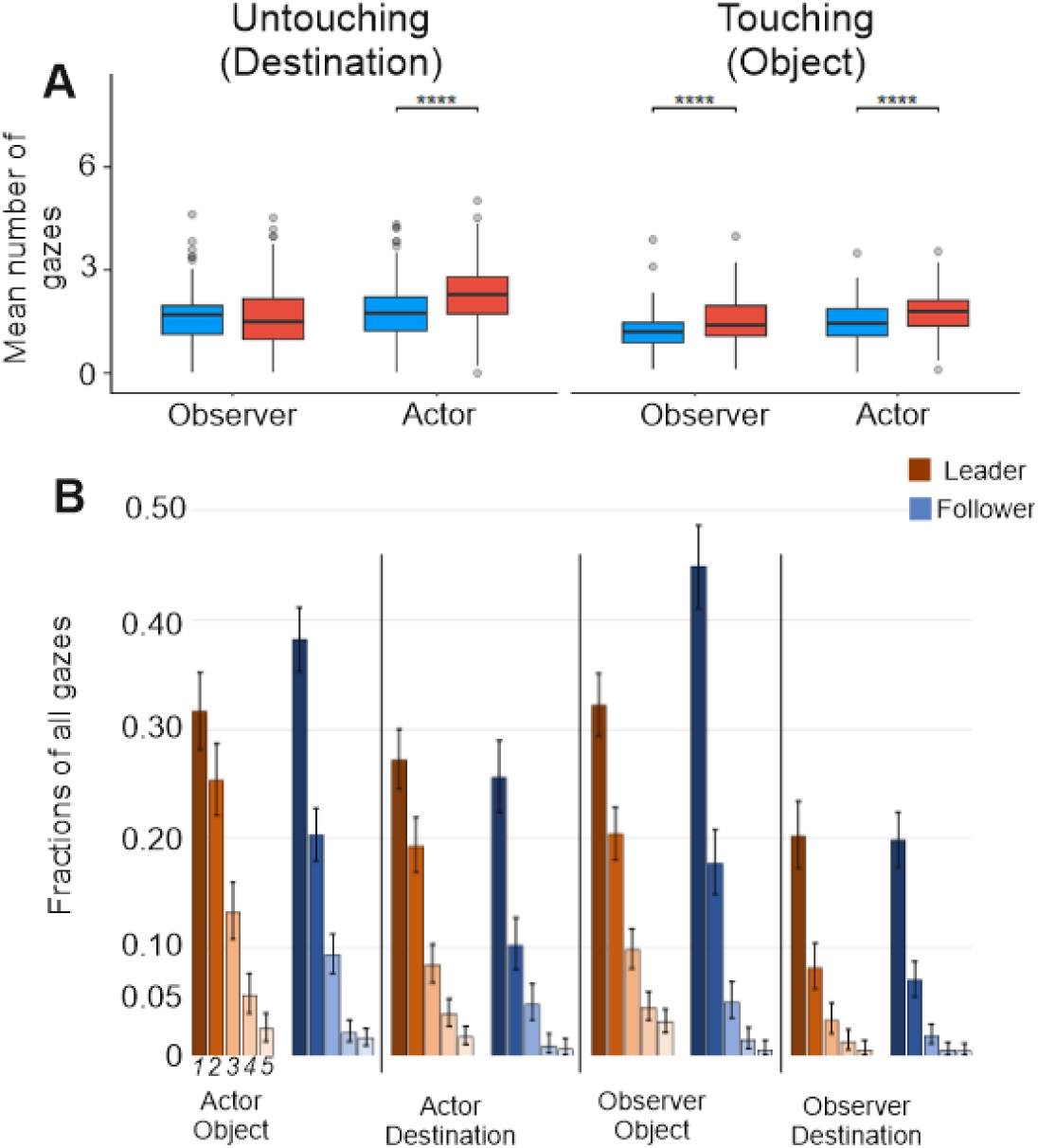
A) Fixation counts for the three-way interaction between Leader, Actor, and Touching factors (**** p<0.0001). B) Single versus multiple fixations, Bars 1 to 5 represents how often a certain fixation has happened. We use a 95% confidence interval calculated for proportions, where the Clopper-Pearson (exact) method for the Binomial distribution was used.

Together, these results indicate that Leaders, compared to Followers, looked at the object that they were moving and its destination more frequently, indicating repeated attention and higher saliency (Irwin, 2013). However, the higher interest of Leaders in the destination of objects vanishes when observing the actions of their partner (i.e., the Follower), showing a shift in resource allocation from following a movement with an already-known outcome to planning their own next movement. One should note that the destination of an object moved back by a Follower, after the Leader’s move, was already clear (for more details about the paradigm, check the method section). More detailed information about the underlying distributions is provided in Figure 7B.

### Fixation Durations

We first focused on fixation length (Eq. 3) and investigated the main effects and interactions via hypothesis testing (Figure 8, for the full results, see Table 2 in the Supplement). The results showed that, when acting (𝑚𝑒𝑑𝑖𝑎𝑛 = 720 𝑚𝑠 ± 570) compared to observing (𝑚𝑒𝑑𝑖𝑎𝑛 = 653 𝑚𝑠 ± 571), participants looked at the target (object as well as destination) significantly longer (*H_0_: Activity* = 0, *Estimate ± SE* = 0.18 ± 0.03, *CI*= [0.12, 0.25], ***p.p.* = 0.00, *BF_01_* =0.00**).

**Figure 8.**
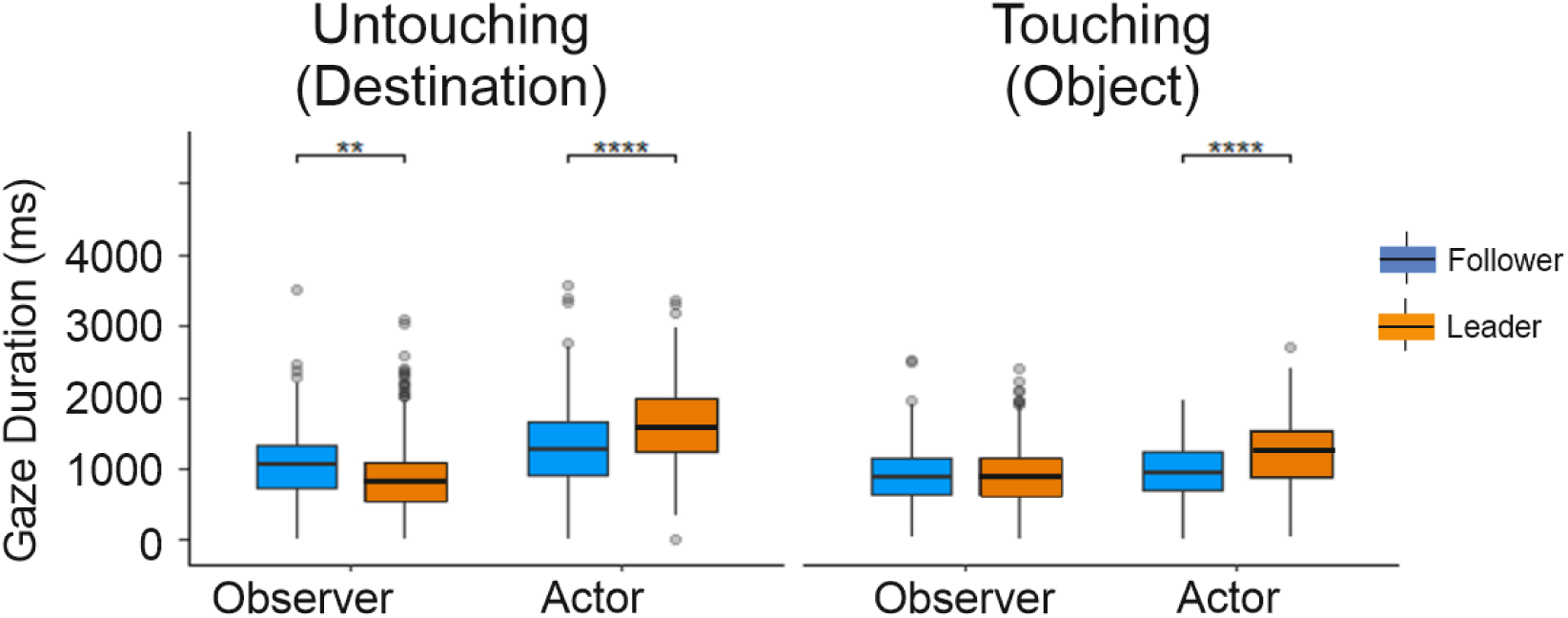
Fixation durations for the three-way interaction between Leader, Actor, and Touching factors. (** p<0.01, **** p<0.0001).

**Table 2.** Hierarchical Bayesian generalized linear modelling results for the fixation duration (Eq. 3 in the main text)

| Hypothesis | Estimate (SE) | 95% CI | BF<sub>01</sub> | Post Prob |
| --- | --- | --- | --- | --- |
| **Activity = 0** | 0.18 ± 0.03 | [0.12, 0.25] | **0.00** | **0.00** |
| Role = 0 | -0.08 ± 0.04 | [-0.15, -0.01] | 4.77 | 0.83 |
| Event = 0 | 0.05 ± 0.03 | [-0.01, 0.11] | **19.66** | **0.95** |
| **Time Window = 0** | 0.14 ± 0.01 | [0.12, 0.15] | **0.00** | **0.00** |
| **Activity \* Role = 0** | 0.20 ± 0.05 | [0.10, 0.29] | **0.01** | **0.01** |
| **Activity \* Event = 0** | -0.19 ± 0.04 | [-0.27, -0.11] | **0.00** | **0.00** |
| Role * Event = 0 | 0.06 ± 0.04 | [-0.02, 0.14] | **16.00** | 0.94 |
| Activity * Role * Event = 0 | -0.01 ± 0.06 | [-0.12, 0.10] | **34.61** | **0.97** |

Note: Significant alternative hypotheses are indicated in bold.
CI: Credential interval.
BF<sub>01</sub>: Bayes Factor comparing the Null hypothesis to the alternative hypothesis. All hypotheses were tested using the brms package (Bürkner 2017). Based on the suggestion of van Doorn et al. (2021), *Bayes factors*(BF) > 3 were considered as significant evidence for the null hypothesis and *BF* < 1/3 for the alternative hypothesis.

The significant interaction between Activity and Event (*H_0_: Activity*Event* = 0, *Estimate ± SE* = -0.19 ± 0.04, *CI*= [-0.27, -0.11], ***p.p.* = 0.00, *BF_01_* =0.00**) revealed that when observing, participants spent more time looking at the actor’s object (i.e., Touching events: 𝑚𝑒𝑑𝑖𝑎𝑛 = 712 𝑚𝑠 ± 628) compared to the actor’s destination (i.e., Untouching events: 𝑚𝑒𝑑𝑖𝑎𝑛 = 570 𝑚𝑠 ± 449). However, when acting, the trend is reversed, meaning participants spent more time looking at the actor’s destination (𝑚𝑒𝑑𝑖𝑎𝑛 = 912 𝑚𝑠 ± 528) rather than the actor’s object (𝑚𝑒𝑑𝑖𝑎𝑛 = 626 𝑚𝑠 ± 688). This interaction reveals that when planning an action to be executed, attention allocation changed qualitatively compared to when observing an action.

Finally, the interaction between Role and Activity was also significant (*H_0_: Activity*Role* = 0, *Estimate ± SE* = 0.20 ± 0.05, *CI*= [0.10, 0.29], ***p.p.* = 0.01, *BF_01_* =0.01**). This result showed that, when being a Leader, participants were more focused on their own objects and destinations (𝑚𝑒𝑑𝑖𝑎𝑛 = 741 𝑚𝑠 ± 685) rather than the Followers (𝑚𝑒𝑑𝑖𝑎𝑛 = 566 𝑚𝑠 ± 465). However, when being a Follower, participants attend to the Leader’s objects and destinations (𝑚𝑒𝑑𝑖𝑎𝑛 = 782 𝑚𝑠 ± 644) more than their own (𝑚𝑒𝑑𝑖𝑎𝑛 = 708 𝑚𝑠 ± 554). These results depict a leader-centered interaction between pairs.

#### Fixation latencies

We investigated fixation latency (Eq. 4) to understand cognitive processes related to planning actions, which consist of two parts: moving an object (i.e., Touching events) and placing it at a destination (i.e., Untouching events). Latency here refers to the time between a fixation on an object (or destination) and the touching (or untouching) event.

First, we checked the main effects and interactions via hypothesis testing (for the full results, see Table 3 in the Supplement). The results (*H_0_: Touching* = 0, *Estimate ± SE* = -0.12 ± 0.03, *CI*= [-0.18, -0.06], ***p.p.* = 0.04, *BF_01_* =0.04**) showed (Figure 9) that participants looked at the destination (𝑚𝑒𝑑𝑖𝑎𝑛 = 2.59 𝑠 ± 5.48) significantly earlier than the object itself (𝑚𝑒𝑑𝑖𝑎𝑛 = 1.95 𝑠 ± 3.2). The significant main effect of Role (*H_0_: Role* = 0, *Estimate ± SE* = 0.15 ± 0.03, *CI*= [0.09, 0.22], ***p.p.* = 0.00, *BF_01_* =0.00**) further showed that fixation latencies when following (𝑚𝑒𝑑𝑖𝑎𝑛 = 1.85 𝑠 ± 4.38), compared to when leading (𝑚𝑒𝑑𝑖𝑎𝑛 = 2.48 𝑠 ± 4.15), were significantly shorter.

**Figure 9.**
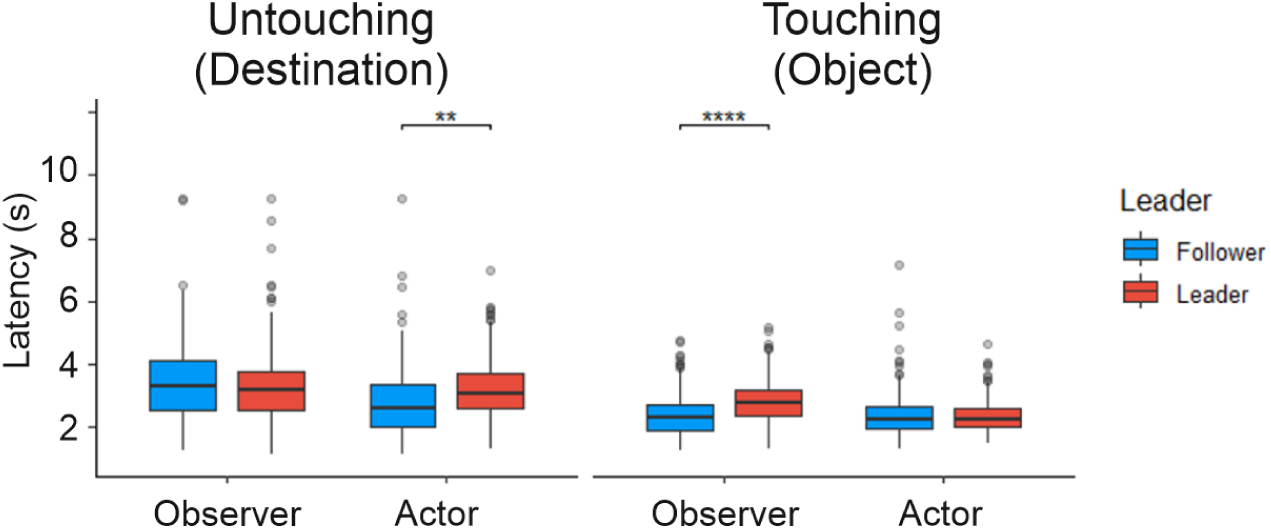
Fixation latencies under different conditions (** p<0.01, **** p<0.0001)

**Table 3.** Hierarchical Bayesian generalized linear modelling results for the fixation latency (Eq. 4 in the main text)

| Hypothesis | Estimate (SE) | 95% CI | Evid Ratio | Post Prob |
| --- | --- | --- | --- | --- |
| Activity = 0 | -0.07 ± 0.03 | [-0.14, 0.00] | **3.12** | 0.76 |
| Role = 0 | 0.15 ± 0.03 | [0.09, 0.22] | **0.00** | **0.00** |
| Event = 0 | -0.12 ± 0.03 | [-0.18, -0.05] | **0.02** | **0.02** |
| **Time Window = 0** | 0.29 ± 0.01 | [0.28, 0.31] | **0.00** | **0.00** |
| Activity * Role = 0 | -0.04 ± 0.04 | [-0.12, 0.05] | **14.55** | 0.94 |
| **Activity \* Event = 0** | 0.14 ± 0.04 | [0.06, 0.22] | **0.12** | 0.11 |
| **Role \* Event = 0** | 0.12 ± 0.04 | [0.04, 0.20] | **0.35** | 0.26 |
| Activity * Role * Event = 0 | -0.13 ± 0.06 | [-0.24, -0.02] | 1.34 | 0.57 |

Note: Significant alternative hypotheses are indicated in bold.
CI: Credential interval.
BF<sub>01</sub>: Bayes Factor comparing the Null hypothesis to the alternative hypothesis. All hypotheses were tested using the hypothesis package from brms (Bürkner 2017). Based on the suggestion of van Doorn et al. (2021), *Bayes factors*(BF) > 3 were considered as significant evidence for the null hypothesis and BF < 1/3 for the alternative hypothesis.

Finally, the two-way interaction between Activity and Event (Figure 9) (*H_0_: Activity*Event* = 0, *Estimate ± SE* = 0.14 ± 0.04, *CI*= [0.06, 0.22], *p.p.* = 0.11, ***BF_01_* =0.12**) and Role and Event (*H_0_: Role*Event* = 0, *Estimate ± SE* = 0.12 ± 0.04, *CI*= [0.04, 0.20], *p.p.* = 0.26, ***BF_01_* =0.32**) were significant. Post hoc tests revealed that when leading compared to following, participants looked at the object to be moved by their partner and the destination of their object significantly earlier. One can understand these results if one notices that the significantly longer fixation latencies for the Leader (𝑚𝑒𝑑𝑖𝑎𝑛 = 2.74 𝑠 ± 3.64) compared to the Follower (𝑚𝑒𝑑𝑖𝑎𝑛 = 1.50 𝑠 ± 3.32) in looking at the object that is moved by the partner indicate that the Leader has advanced information about which object is going to be touched by the Follower. However, Leaders (𝑚𝑒𝑑𝑖𝑎𝑛 = 2.77 𝑠 ± 4.93), compared to Followers (𝑚𝑒𝑑𝑖𝑎𝑛 = 2.04 𝑠 ± 5.14), look significantly earlier at the destination of the object that they are moving. This result shows that Leaders allocate more resources to planning their actions fully from the beginning compared to followers (i.e., both the object that should be moved and its destination).

#### Sequences

Finally, we asked whether participants produced sequential combinations of fixations on objects and destinations. Such patterns are particularly informative for understanding decision processes that unfold after a participant has fixated on the object they will soon manipulate—coded on the abscissa in Figure 10 as a leading “0”—or on the destination where that object will be placed—coded as “1.” To examine this, we analyzed potentially relevant 3- and 4-fixation sequences beginning with 0 or 1. For example, a sequence coded as “0.1.0” denotes a look at one’s own object (“0”), followed by a look at its destination (“1”), and then a return to the object (“0”); the corresponding bar in the figure indicates the proportion of such sequences. Indices “2” and “3” refer to the partner’s subsequent targets (object and destination). Figure 10 plots the frequencies of several such sequences, while the final columns provide baseline values derived from gaze sequences directed at objects not involved in the manipulation during the given episode (bars labeled “xyx” and “xyz” in panels A and B). Error bars represent binomial confidence intervals, computed using the Clopper-

**Figure 10.**
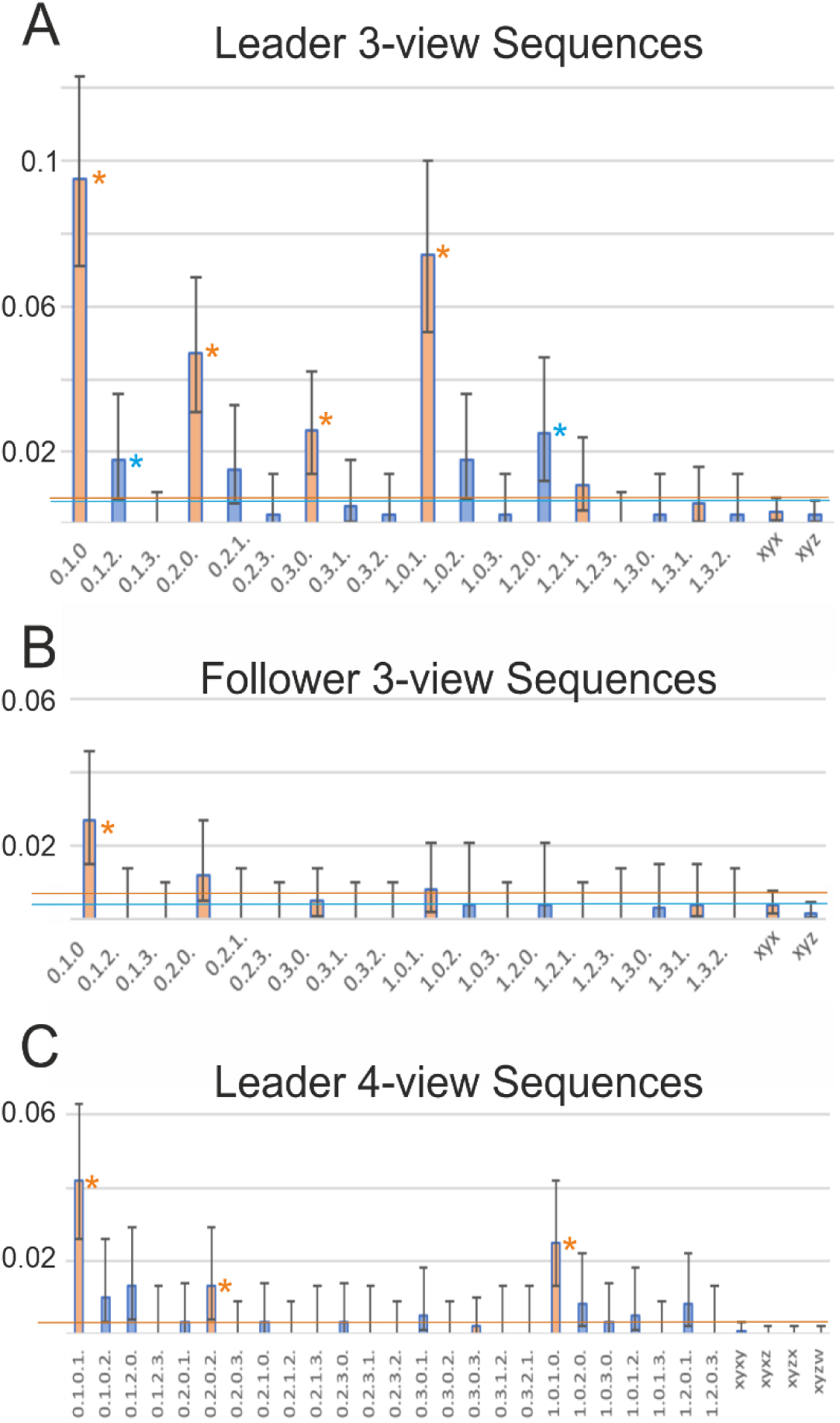
Sequences, A,B) 3-view sequences, C) 4-view sequences only for the Leader. Numerals mean: 0=own object, 1=own destination, 2=the other’s object, 3=the other’s destination.

Pearson (exact) method. The horizontal lines, color-coded by sequence structure, indicate the confidence intervals of the baseline across the diagram. A sequence can therefore be considered above chance when its bar, including the lower confidence bound, lies entirely above the corresponding baseline line.

Leaders (panels A, C) indeed performed several sequences significantly above chance. All combinations of 010 and 101, as well as their 4-fixation counterparts 1010 and 0101, were overrepresented. (Note that the 3-fixation sequences are naturally contained within their 4-fixation extensions.) Sequences 020 and 0202 were also prevalent, indicating alternations between the Leader’s own object and the partner’s-folower’s object. In addition, a small overrepresentation was found for 030 (Leader’s object– partner’s-follower’s destination–Leader’s object), although its 4-fixation counterpart 0303 was rare.

Mixed sequences involving three different entities (x, y, z) were observed only for the combination of 0 = own object, 1 = own destination, and 2 = partner’s object. Specifically, the sequences 012, 021, 102, and 120 occurred, with two (012 and 120) just above chance and the other two just below. This pattern suggests at least a tendency for Leaders to integrate their own object and destination with the partner’s object in triadic sequences.

For the Follower (B), the first four orange bars resembled a compressed version of the corresponding bar for the Leader (A). However, only the 3-view sequence 010 was significantly above chance. No 4- view sequences reached significance for the Follower, so this plot is omitted.

Taken together, these findings show that Leaders often linked their own objects and destinations in extended alternations, sometimes incorporating the partner’s objects or destinations as well.

Followers, by contrast, showed much simpler sequential patterns, largely limited to alternations between their own object and its destination.

## Discussion

This study investigated how gaze and action unfold in a continuous, role-based interaction. Unlike many laboratory tasks with artificial cues or parallel cooperation, our “do–undo” game required alternating Leader and Follower roles. In this setting, the Follower’s planning necessarily depended on the Leader’s behavior, while the Leader retained more autonomy but could still anticipate the Follower’s next moves. Our combination of dual eye tracking, multi-camera motion capture, and touch sensors enabled us to segment cooperative action into precise temporal intervals and to demonstrate not only the well-established precedence of gaze before action, but also striking role differences in how Leaders and Followers plan and monitor their behavior.

Clear differences also emerged between acting and observing. Leaders made more fixations when acting than when observing, a pattern not seen in the Follower. At one level, this reflects the task structure: Leaders were responsible for initiating each move and therefore had to invest more visual planning, whereas Followers only needed to identify what had to be undone. What is informative, however, is that the gaze data reveal how this division of labor translated into attentional priorities. Leaders devoted most of their fixations to planning their own actions, with comparatively little monitoring of the Follower, while Followers focused selectively on the Leader’s object choices, largely ignoring destinations. These asymmetries illustrate how role assignment shapes not only motor behavior but also the allocation of visual attention in joint action.

Further asymmetries between Leader and Follower were evident in fixation latencies. That Leaders look-ahead to both the object and the destination was, to some extent, imposed by the task structure: the Leader had to initiate each new sequence, whereas the Follower could only act in response. What is noteworthy, however, is how clearly this asymmetry manifested in the gaze data, revealing a consistent temporal gap between proactive planning in Leaders and reactive adjustment in Followers (see Figure 6A). This pattern resonates with broader accounts of proactive versus reactive control, and highlights how role assignment in joint action shapes the timing and content of visual planning.

Fixation duration also proved informative, echoing findings by van der Laan et al. (2015). Actors showed reliably longer fixations than observers, reflecting the greater attentional investment required for action planning. Importantly, the interaction between activity and event revealed a systematic shift in visual priorities: during action, participants fixated longer on the destination, whereas during observation, they focused more on the object to be moved. This dissociation demonstrates how role assignment shapes the functional allocation of gaze — either to guide one’s own unfolding movement or to track the co-player’s choice of object. The pattern aligns with previous work showing that in pick-and-place actions, people fixate on the object until it is lifted and then shift their gaze to the destination until release (Lavoie et al., 2018). Consistent with this, predictive gazes have also been documented in observers, who tend to look ahead to the actor’s goal rather than simply tracking the hand (Flanagan et al., 2013). Our study complements these findings by disentangling how such predictive gaze patterns differ between acting and observing in a cooperative, role-based task.

Leaders often produced alternating sequences of fixations between object and destination (Figure 10), a hallmark of planning-related search behavior. Look-ahead fixations of this type have been described before (Sullivan et al., 2021; Mennie et al., 2007; Pelz & Canosa, 2001). What our data add is evidence that such fixations frequently unfolded as extended back-and-forth sequences, linking objects and destinations in longer planning chains. This suggests that Leaders were not merely preparing the next immediate move but engaged in multi-step planning, a form of anticipatory gaze behavior not systematically documented in earlier work.

Our analyses further show that Leaders’ predictive eye movements extended beyond the next immediate action. As illustrated in Figure 6G, gaze peaks in intervals 4 and 5 revealed that even while waiting to act, Leaders frequently inspected potential objects the Follower might manipulate. This suggests that Leaders sometimes simulated not only their own upcoming moves but also the likely responses of their partner. Overrepresented sequences such as “own object–follower’s object–own object” or “own object–follower’s location–own object” point to a form of dual planning in which Leaders anticipated both their own and the Follower’s future actions. Such role-crossing predictions highlight the cognitive depth of coordination in this task.

Followers also displayed anticipatory gaze. Within ∼400 ms after the Leader’s hand movement onset, they had already fixated the Leader’s intended target object (Figure 6). This rapid anticipation indicates that Followers did not simply trail the Leader’s hand but actively predicted the unfolding action. In this way, coordination in the task reflected a reciprocal prediction process, with Leaders sometimes anticipating the Follower’s responses and Followers anticipating the Leader’s choices.

Another striking observation concerns the “tails” following the main gaze peaks shown in Figure 6, visible for both acting Leaders and, even more, observing Followers. Many of these fixations were directed to locations where an object had just been removed, effectively gazes into the past. We interpret these retrospective looks as part of a memorization process (O’Regan, 1992; Johansson & Johansson, 2014; Foerster, 2019). For Followers, remembering the pre-action state was essential to perform the undo operation. For Leaders, such retrospective checks may have served to verify the configuration and to monitor whether the Follower subsequently executed the correct undo action.

In sum, this study highlights how Leader and Follower roles elicit distinct decision-making and planning strategies in a continuous, naturalistic setting. Leaders displayed extended sequences of proactive fixations, sometimes spanning several steps and even anticipating potential moves of their partner. Followers, meanwhile, rapidly identified the Leader’s intended targets and combined predictive with retrospective gaze patterns to support the undo task. Both roles also exhibited memory-related looks to previously occupied locations. Together, these findings reveal that joint action relies on a dynamic interplay of proactive planning, reciprocal prediction, and retrospective checking—processes that extend beyond the immediate next move and underscore the complexity of everyday human coordination.

## Acknowledgements

This publication was funded by the Deutsche Forschungsgemeinschaft (DFG, German Research Foundation) - Project-ID 454648639 - SFB 1528, "Cognition of Interaction", Project B01 and by Lower Saxony Ministerium für Wissenschaft und Kultur (MWK), Project: Kognitiv und Emphatisch Intelligente Kollaborierende Roboter (KEIKO), TP6.

## Compliance and declarations

- The work is original, not published previously, and is not under consideration elsewhere.
- All authors have approved the manuscript and agree with its submission to the Journal of Vision.
- The study was conducted in accordance with the Declaration of Helsinki and approved by the Ethics Committee of the University of Göttingen, Psychology (Registration Nr: 294).
- Data, materials, and analysis details are available upon reasonable request.
- The authors declare that there are no known competing interests.

## Supplement for

This supplement provides additional information concerning the computer vision algorithms used to record experimental data as well as results and diagrams for different additional aspects including the statistical-modeling based analysis.

### Computer vision - Ray tracing

The algorithm consists of two main parts: ray casting and tracing. By combining the head pose data and the eye tracker – camera setup transformation matrix, the fixation point is converted into a ray in the 3D space, where its origin is the origin of the scene camera, and the direction vector is calculated by connecting the origin and the fixation point. Mathematically, a ray *r*, defined as 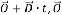 represents the origin of the ray, and it is the origin of the scene camera of the eye tracker. D is the direction vector, which connects the origin and the gaze fixation point on the scene camera’s image plane, 𝑡 is a positive scalar that extends the ray from its origin towards infinity. The tracing algorithm is essentially a collision detection between the casted ray and the 3D AABBs, and it operates by treating each axis individually and then combining their results. In each dimension, a 3D AABB is defined by two lines: one with minimum coordinates and one with maximum coordinates. The intersection area of these lines in 3D space forms the AABB. Each line 𝑙_𝑛,_can be expressed as 𝑌 = 𝐵_𝑛,𝑚_, where 𝑛 is the index of the dimension and 𝑚 can be either "*max*" or "*min*". The solution of the following equation indicates where does ray *r* hits one of these lines: 𝑂_𝑛_ + 𝑡_𝑛,_ · 𝐷_𝑛_ = 𝐵_𝑛,𝑚_. The result 𝑡_𝑛,_give the ranges where the ray *r* hits the boundary lines of the AABB in each dimension by illustrating the extent of the direction vector *D*. In the case where *r* hits the AABB, there will be an overlap range across the ranges of all dimensions, that is: 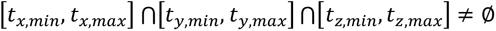 Moreover, there will be a pair of 𝑡_𝑛,𝑚_ that define the range of the ray which penetrates through the AABB, 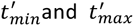 can be used to annotate them. Then a point 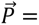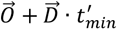 is the point where the ray hits the surface of the AABB in the first place. In contrast, when the ray *r* does not hit the AABB, there is no overlap of t*_n,m_* across all dimensions. By executing the ray tracing algorithm with each 3D AABB, the location data can be extracted. If the ray intersects both an object and a hand, the hand is prioritized as the output.

**Figure 11.**
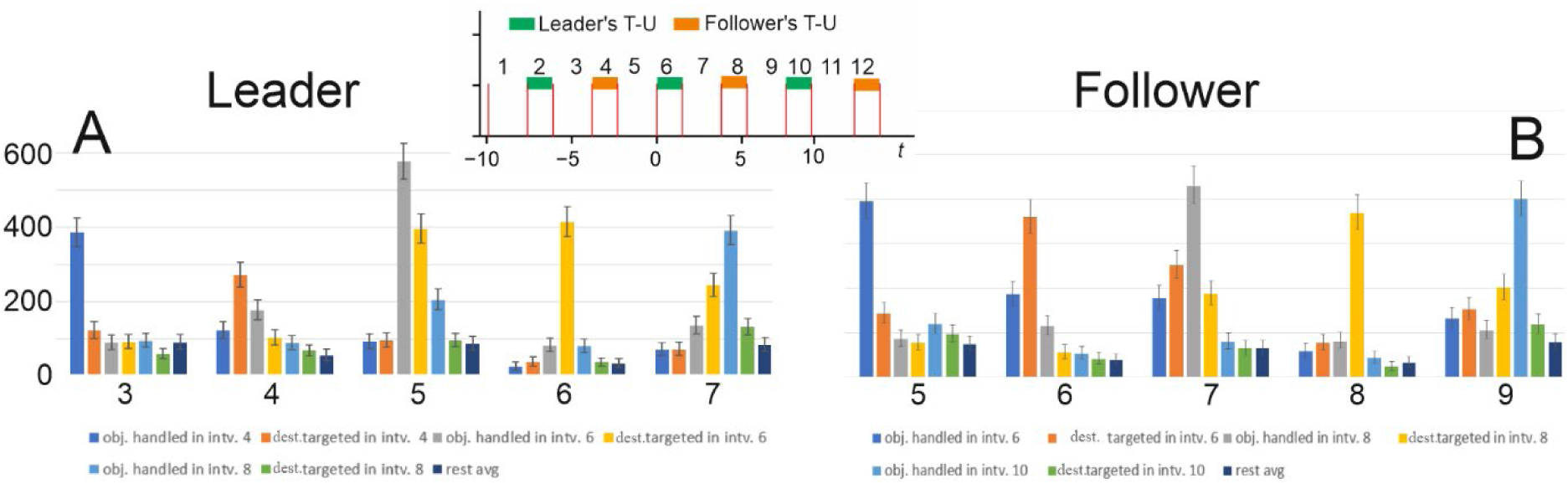
Objects and destinations per interval. Intervals (abscissa) are numbered as shown in the inset above.

### Fixation distribution in consecutive T-U and U-T intervals

In Figure 1 we analyze sequences of time intervals (T-U and U-T), considering at which objects/destinations Leader and Follower have looked in those different intervals. In a nutshell: This diagram shows, quite expectedly, that the different “aspects of interest” are indeed looked at one after the other.

A total of 500 sample-intervals has been analyzed to obtain a histogram for each interval. Thus, histograms show the number of fixations per 500 intervals. Error bars represent 95% confidence intervals calculated for Poisson distributions.

The Leader has finished their action in interval 2 and begins considering their next action but before they look quite often at what the Follower does. Hence, fixations at the Follower’s object (blue bar, left) are common in interval 3 (compare also to Figure 6 G in the main text). Note that these fixations occur before the Follower has actually touched their object, which happens only at the start of 4. During this time the Leader shifts gaze to the targeted destination of the Follower (orange bar). However, the Leader now also begins to consider, which object to manipulate next (gray bar). As expected, this becomes prevalent in interval 5, but here the Leader also starts to search for the placing destination (yellow bar). During execution of the Leader’s action (intv. 6) fixation on the destination is dominant. Note that interval 7 is equivalent to 3 but not identical. This is due to the fact that intervals follow each other to define an episode and the later coming intervals are conditional on the earlier ones, leading to similar but not identical results for equivalent intervals. Hence for 7 the light blue bar in 7 corresponds to the dark blue one in 3 (same for green and orange). Indeed, in interval 7 the fixations at the next object of the Follower again begin to dominate (as in 3), but here the Leader still keeps looking at their chosen destination (yellow), too.

For the Follower a quite similar pattern emerges. However, in interval 6 and 7 the dark blue and orange bar are larger for the Follower than the equivalent counterparts for the Leader (which are found in intervals 4 and 5). For both of them, these two bars represent “what the other does” and, evidently, this is more relevant for the Follower than for the Leader.

### Additional material about the statistical modelling-based analysis

Here we provide additional material concerning the statistical analyzes of the fixation patterns (length, number, latency). In all three cases the statistical model converged correctly without any divergent transitions (all *Rhat* = 1) as described in the main text. Further, bulk and tail effective sample sizes for main effects and interactions were above 4000, showing that model predictions were reliable. Furthermore, the model accurately captured the observed data distribution.

### Fixation counts

In Table 1, we provide details on hypothesis testing for fixation counts, i.e., the number of fixations as modeled by Equation 2, with main factors of activity (Actor vs. Observer), role (Leader vs. Follower), and event (Touching vs. Untouching events). For details on the model, please refer to the main text.

### Fixation duration

In Table 2, we provide details on hypothesis testing for fixation duration, i.e., the cumulative duration of fixations, as modeled via Equation 3. For details on the models, please refer to the main text.

### Fixation latency

In the same way as for the other two aspects above, in Table 3, we provide details on hypothesis testing on fixation latency, i.e., how far in advance of a predefined event a fixation is elicited as modeled via Equation 4. For details of the model, please refer to the main text.

## Footnotes

1 Note: only direct repetitions were forbidden by the rules of the game, but a repetition by manipulating an object in interval 2 and then again at 10 would be permitted.

## Notes

### Competing Interest Statement

The authors have declared no competing interest.

